# *In vitro* assessment of an engineered tBID-based safety switch system in human T lymphocytes

**DOI:** 10.1101/2021.01.27.428458

**Authors:** Jiamiao Lu, Patrick Collins, Ki Jeong Lee, Chi-Ming Li, Songli Wang

**Author notes:** Equal contribution (Jiamiao Lu, Patrick Collins). **Correspondence Authors:** Chi-Ming Kevin Li PhD; MSc, Songli Wang MD; PhD, 1120 Veteran Blvd, ASF1-2, South San Francisco, CA94080.

## Abstract

Cell therapy as a promising therapeutic modality to treat cancer has been intensively studied for decades. However, the clinical trials have indicated that patients under T cell therapy may develop severe cytokine release syndrome resulting in hospitalization or even death. Furthermore, genetic modifications to promote proliferation and persistence of T cells could result in high numbers of long-lived engineered cells in patients after treatment. In this study we developed a novel tBID-based safety switch regulated through a small molecule inducible on switch to control undesired toxicity or ablate engineered cells as needed. We compared several tBID-based safety switch constructs with the clinically validated safety switches, HSV-TK (human herpes simplex virus thymidine kinase) and iCasp9 (inducible Caspase 9), few tBID-based safety switch constructs were systematically tested and investigated *in vitro* in Jurkat or human primary T cells. We demonstrated that our novel tBID based safety switch, with optimization, was able to eliminate up to ~90% of transduced human primary T cells within 72hr after activation, providing an alternative switch system to manage safety concerns for cell therapy.

## BACKGROUND

Over the last few decades cell therapies have opened opportunities to new treatments for cancers, autoimmune disorders, and inherited diseases. These treatments often involve transplantation of genetically-engineered stem cells or specific cell types of interest. However, the potentials of the exogenously infused cells to malignant transformation and the toxicities of the transgene-expression products have led to the emergence of safety concerns. Numerous safety switch systems have been incorporated and developed to mitigate undesired toxicities in the donor cells, resulting in selective depletion when severe toxicities, for examples cytokine release syndromes, arise.

A safety switch derived from the human herpes simplex virus thymidine kinase type 1 gene, HSV-TK, has been clinically validated in allogenic donor T cell infusion to treat tumor and Epstein Barr virus (EBV) lymphoproliferative disease after hemopoietic stem cell transplantation (1–3). The killing activity of HSV-TK is activated by an antiviral prodrug, Ganciclovir (GCV), which hinders the administration of GCV for cytomegalovirus infections. In recent years a robust fast killing safety switch system, inducible Caspase 9 (iCasp9), has emerged (4,5). In this system, the human Caspase 9 protein lacking the caspase activation and recruitment (CARD) domain is fused to a human FK506-binding protein (FKBP) derived drug-binding domain. Without activation the Caspase 9 fusion protein stays in monomer form. By binding with a chemical induction of dimerization (CID) drug, AP1903, the Caspase 9 fusion protein dimerizes and in turn activates the cellular apoptosis events. The feasibility of iCasp9 has been tested clinically in the patients who received haploidentical hematopoietic stem cell transplantation, in which the genetically-engineered donor T cells decreased by more than 90% within 30 minutes after CID infusion (5). The efficacy of iCasp9 in controlling serious adverse events caused by genetically-engineered chimeric antigen receptor (CAR) T-cell therapy was also validated in phase 1 clinical trials (6). However, while iCasp9 responded quickly in 30 minutes upon induction, it could take a few days for the HSV-TK system to show effects.

As one of pro-apoptotic BCL-2 family members, BID functions as a BH3 domain only death agonist (7). The cytosol-localized full-length BID is a substrate of Caspase 8 (8). Cleavage of BID by Caspase 8 releases its pro-apoptotic activity. The truncated BID (tBID) then translocates from cytosol to mitochondria where it induces clustering of mitochondria around nucleus, cytochrome c release, and then loss of mitochondrial membrane potential, leading to cell shrinkage and nuclear condensation (8).

While tBID demonstrates many of the properties of an ideal kill switch, expression of the protein itself would lead to cell death and so it needs to be coupled to some sort of a molecular switch. The ideal switch would result in no background expression of tBID until induced by a small molecule. Ideally, the inducer would be commercially available, clinically validated, inexpensive, well-characterized compound that can pass through both the blood-brain barrier (BBB) and the blood-retinal barrier (BRB). Trimethoprim (TMP) is a frequently used antibiotic that binds to bacterial dihydrofolate reductase (DHFR) with 1000X selectivity compared to human DHFR. Multiple *E. coli* dihydrofolate reductase (ecDHFR) destabilizing domain mutants have been engineered for construction of the destabilizing domains (9). By fusing such a destabilizing domain to the other domain or protein of interest, the entire fusion protein should be unstable and rapidly degraded. Addition of TMP stabilizes the destabilizing domain in a rapid, reversible and dose-dependent fashion (10,11), suggesting that cell death mediated by TMP administration may be a practical method to manage or attenuate safety concerns in cell therapy.

In this study, we took the advantage of the ecDHFR destabilizing domain system fused with tBID to develop an alternative safety switch system that can not only gradually but also thoroughly deplete engineered donor cells to expand the options of safety switch systems. The safety switch candidates were constructed by fusing tBID with the mutants of the ecDHFR destabilizing domain at either N- or C-termini. The potency of these tBID based safety switches were validated and assessed *in vitro* with or without TMP in both human Jurkat and primary pan-T cells.

## MATERIALS AND METHODS

### Vector construction and cloning

The iCasp9, NDHFR-tBID, HSV-TK, and tBID-CDHFR fragments were synthesized by GENEWIZ. These fragments were then cloned into in house lentiviral vectors. The M97A/D98A mutations were introduced into NDHFR-tBID by using In-Fusion site directed mutagenesis kit with forward primer 5’CCAGGTCGGGGACAGCGCGGCGCGTAGCATCCCTCCGGGC3’ and reverse primer 5’GCCCGGAGGGATGCTACGCGCCGCGCTGTCCCCGACCTGG3’.

### Cell culture and transduction

The Jurkat cells (ordered from ATCC) were cultured with RPMI1640 medium supplemented with 10% FBS (Fetal Bovine Serum) and 100 μg/ml streptomycin. On the day performing transduction, lentiviral vector at desired MOI was added to 1ml of 2xE+05 cells/ml in 15ml Falcon tube in PRMI1640 medium with 10% FBS and 5 μg /ml polybrene then followed by gentle agitation. Spinoculation was performed by centrifugation of the tubes at 1200g for 2hr at 32 °C. Supernatant was aspirated off, resuspended cells with 2ml PRMI1640 medium with 10% FBS, placed the cells in 12 well plate and incubated the plate in incubator at 37°C.

The frozen human Pan-T cells (ordered from AllCells, LLC.) were thawed and cultured with X vivo 15 medium, 5% heat-inactivated AG serum-Sigma, 1XPen/strap, anti-CD3/anti-CD28 Dynabeads (at 1:1 cells: beads ratio) at 1xE+06 cells/ml per well in 24-well plate on day 0. Performed CD3 staining on day 0 to confirm T cell purity. Transduction was performed on day 3. First added fresh medium with IL-2 100U/ml to make the cell concentration at 0.5xE+06 cells/ml. Place 1ml of cell suspension into each 15ml falcon tube. Then added virus at desired MOI and 8μg/ml polybrene. Span the tubes at 1800rpm (1000g) for 45min at 32 °C. Finally removed virus, replaced the medium at 0.5E6 cells /ml with IL-2 100U/ml, and culture cells at 37 °C tissue culture incubator. Changed medium when needed.

### Protein L magnetic bead purification of lentivirus transduced human pan-T cells

1xE+06 transduced human pan-T cells were counted and washed with cold PBS on day 3 post lentivirus transduction. The cells were then resuspended in 1ml cold PBS with 0.5% BSA and 0.05% Tween-20. 100μl of Protein L magnetic beads (Pierce Protein L Magnetic Beads, Cat# 88850) were first washed with 1ml cold PBS with 0.5% BSA and 0.05% Tween-20 by gentle vortexing for 1 min. Then collected the beads by using magnetic stand and removed supernatant. Then added the resuspended transduced human pan-T cells into the tube with washed beads. Incubate at 4°C with mixing for 1~2hr. Collected enriched cells by magnetic stand and removed supernatant. Wash the cells with 1ml cold PBS with 0.5% BSA and 0.05% Tween-20 twice by using magnetic stand. Removed the wash buffer after the final wash. Resuspended the final bead-cell pellet with fresh X vivo 15 with IL-2 100U/ml and incubated in cell culture flask at 37°C. Beads fell off after incubation and could be removed by magnetic stand before next analysis.

### Flow cytometry and bead-based cell counting

On day 3 post lentivirus transduction in human pan-T cells and Jurkat cells, prepared cell suspension for each group at 1xE+06 cells/ml, and put 10ml cell suspension in T75 flask (two flasks for each transduced construct). Treated one flask of each transduced construct group with DMSO and the other flask with 100nM TMP in PRMI1640 medium with 10% FBS for Jurkat cells and X vivo 15 medium, 5% heat-inactivated AG serum-Sigma, 1XPen/strap, andIL-2 100U/ml for human pan-T. The killing efficacy was quantified by flow cytometry combined with BD liquid counting beads (BD Pharmingen Cat# 335925) on day 1, day 2, and day 3 post treatment. On the day of flow analysis, first took 1ml cell culture into 1.5ml Eppendorf tube from each flask, added 50ul BD liquid counting beads to each tube and span. Then remove medium and added 100μl BD staining buffer (BD Pharmingen Cat# 554657) containing Pierce Recombinant Protein L, Biotinylated (Thermo Scientific Cat# 29997) at 1:100 ratio. Incubated for 30min at 4°C, then washed cells twice with 1ml PBS. Removed PBS and added 100μl of BD staining buffer with PE Streptavidin (BD Pharmingen Cat# 554061) as secondary (1:10 dilution) and incubate for 30min at 4°C, then washed twice with PBS. Then removed buffer and resuspended cells and beads in BD staining buffer with Annexin V-Cy5 (Abcam Cat# ab14150) incubated for 10min in dark at room temperature. Span and removed buffer. Then fixed cells with 1% paraformaldehyde in BD staining buffer for 60min at room temperature. Then washed cells twice with 500μl BD staining buffer then resuspend in 200ul BD staining buffer for flow cytometry analysis. For Figure 1D, the elimination rate of GFP positive cells was calculated based on the following formula:

**Figure 1.**
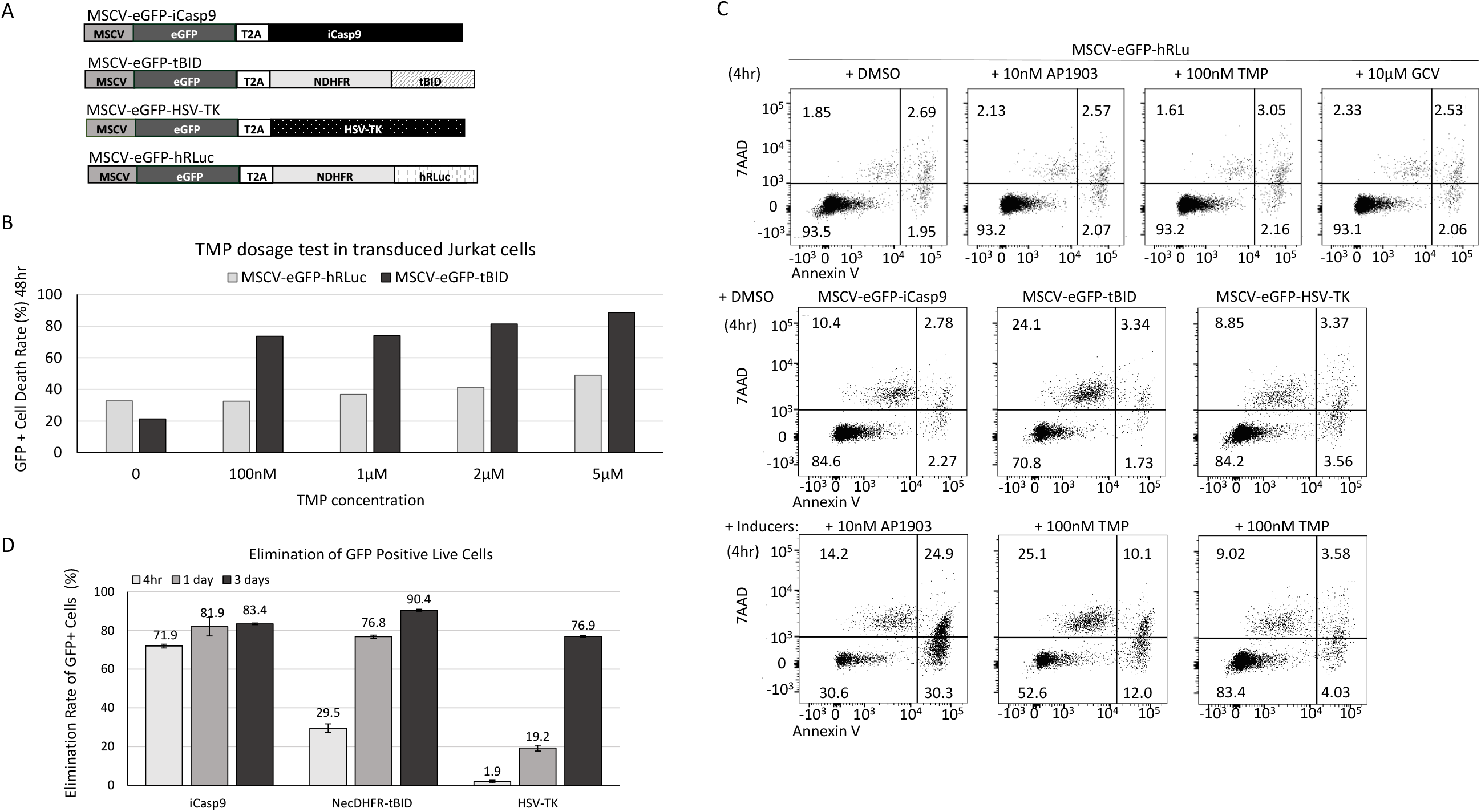
The killing efficacy of NDHFR-tBID safety switch in Jurkat cells. A) Schematic diagrams of the lentivirus vectors used to transduce Jurkat cells. B) Dosage test of TMP for activating tBID safety switch in Jurkat cells. The Jurkat cells transduced by *MSCV-eGFP-hRLuc* lentivirus or *MSCV-eGFP-tBID* lentivirus were treated with TMP at numerous concentrations for two days. The viability of transduced Jurkat cells were quantified through flow cytometry analysis after staining with Annexin V and SYTOX Blue. The percentage of dead cells was then plotted. The X axis indicates the concentration of TMP. The Y axis indicates the rate of cell death (%). C) Flow cytometry analysis of Jurkat cells transduced with lentivirus vectors described in A at 4hr post treatment. All lentivirus vectors described in A expressed puromycin resistance gene. Transduced Jurkat cells were selected with 0.5 μg/ml puromycin for 14 days before treatment with DMSO, AP1903, TMP and GCV. Cells were stained by viability dye 7AAD and Annexin V at 4hr post treatment. D) The elimination rate of GFP positive Jurkat cells transduced with lentivirus vectors described in A at various time points post treatment with DMSO, AP1903, TMP and GCV. To calculate the elimination rate of GFP positive cells transduced with each lentivirus vector, the percentage of GFP positive viable cells in the DMSO control compound treated group of each lentivirus vector transduced cohort was first used to subtract the percentage of GFP positive viable cells in the AP1903, GCV, or TMP treated group of the same cohort at the same time point, then divided by the percentage of GFP positive viable cells in the DMSO control compound treated group, and then times 100%.

Elimination rate of GFP positive cells (%) =

(% of DMSO treated GFP positive live cells - % of AP1903/GCV/TMP treated GFP positive live cells) / % of DMSO treated GFP positive live cells x 100%

## RESULTS

### The tBID-based safety switch demonstrated slower but compatible killing efficacy with iCasp9 in Jurkat cells

In order to develop a novel, regulatable safety switch, we incorporated the pro-apoptotic role of tBID in cell death with the destabilizing feature of the mutant ecDHFR destabilizing domain to form the tBID-ecDHFR fusion protein (9). Theoretically, in the absence of TMP, the tBID-ecDHFR fusion protein would be unstable and quickly degraded. However, upon TMP treatment, the fusion protein should become stabilized and allow tBID to induce cell death. In order to test this hypothesis, tBID was fused to the N-terminal ecDHFR destabilizing domain (NDHFR) mutant (R12Y/Y100I) (9) (Figure 1A) because the C-terminal ecDHFR destabilizing domain (CDHFR) mutant (N18T/A19V) had been demonstrated to be leaky in the absence of TMP treatment in previous literatures and our own experiment (Supplementary Figure 1) (9). Since even low levels of tBID fusion protein could lead to undesired basal toxicity, the NDHFR-tBID fusion construct was chosen at this stage for efficacy assessment and subcloned into a lentiviral expression vector containing eGFP and puromycin resistance genes driven by the *MSCV* promoter (Figure 1A). In the meantime, two well-established safety switches, iCasp9 and HSV-TK, and a NDHFR-fused humanized Renilla luciferase (*hRLuc*) were also incorporated in the same lentiviral vectors as the positive and the negative controls, respectively (Figure 1A). Although the dosages of AP1903 and GCV to activate iCasp9 and HSV-TK, respectively, were well established (3–6), the functional dosage of TMP was less-defined and further tested in the Jurkat cells transduced with lentivirus vectors *MSCV-eGFP-hRLuc* and *MSCV-eGFP-tBID* (MOI=10). MOI=10 was chosen since it provided better transduction efficacy when compared with MOI=1 (Supplementary Figure 1A). After two weeks of puromycin selection, the transduced Jurkat cells were treated with TMP at various concentrations (0, 100nM, 1μM, 2μM, and 5μM) for two days. As shown in Figure 1B (Supplementary Figure 2B), 48hr of TMP treatment at 100nM was able to increase cell death from 22.3% (no TMP treatment) to 73.6% in the *MSCV-eGFP-tBID* transduced cells without inducing additional cell death in the control-vector, *MSCV-eGFP-hRLuc*, transduced cells. However, TMP treatment at higher concentration (2μM and 5μM) induced additional cell death in the control group (Figure 1B, Supplementary Figure 2B), indicating potential toxicity at higher concentration. Therefore, the lowest concentration of TMP (100nM) was used in this study to activate the tBID-based safety switch.

To compare the efficacy of these constructs to induce cell death, the packaged lentiviral vectors were transduced into Jurkat cells at MOI=10. After two weeks of puromycin selection, the transduced Jurkat cells were treated with either dimethyl sulfoxide (DMSO) as the vehicle control reagent or 10nM AP1903 to activate iCasp9, 100nM TMP to activate tBID, and 10μM GCV to activate HSV-TK, respectively. To investigate cell viability affected by the treatment, the viability of the positively-transduced cells (GFP positive) was monitored and quantified through flow cytometry analysis after 7-amino-actinomycin D (7AAD, a DNA dye staining late apoptotic cells) and Annexin V (a marker binds to cell surfaces expressing phosphatidylserine and recognizes early apoptotic cells) staining at a serial time-points (4hr, day 1 and day 3). As shown in Figure 1C, at 4hr after different compound treatments, no viability change was detected in the control *MSCV-eGFP-hRLuc*-transduced Jurket cells treated with different small molecules or the DMSO-treated cells transduced by the various lentivirus preps. However, at the same time-point, strong cell death was induced in iCasp9 construct (*MSCV-eGFP-iCasp9*) transduced group, whereas only mild induction in cell death was observed in the cells transduced with *MSCV-eGFP-tBID* virus and very rare induction in cell death was observed in the *MSCV-eGFP-HSV-TK* transduced cells (Figure 1C and 1D).

To further analyze and quantify the flow cytometry results on days 1 and 3, the elimination rate of the GFP-positive cells was calculated (Supplementary Figure 3). As plotted in Figure 1D, acting as a fast-acting safety switch, iCasp9 was able to eliminate 71.9% GFP positive cells at 4hr after being activated by AP1903. On day 1 and day 3 post AP1903 treatment, the activated iCasp9 eliminated 81.1% and 83.4% GFP-positive cells, respectively. On the other hand, the slow acting safety switch, HSV-TK, showed no obvious killing activity at 4hr after GCV treatment. However, it’s killing effects started gradually, with 19.2% elimination rate on day 1 and 76.9% elimination rate on day 3 post GCV treatment. The NDHFR-tBID safety switch demonstrated a moderate killing effect that fell into the range between iCasp9 and HSV-TK. The elimination rates of the NDHFR-tBID engineered Jurkat cells with TMP treatment at 4hr, day 1 and day 3 time points were 29.5%, 76.8% and 90.4%, respectively (Figure 1D).

### NDHFR-tBID activity was attenuated in human pan-T cells

After validating the efficacy of NDHFR-tBID safety switch in the Jurkat cells, a T-cell leukemia cell line, we incorporated the NDHFR-tBID and a control eGFP expression cassette, respectively, into a CAR expression vector under the control of *hEF1α* promoter to compare and assess their killing activities in primary human pan-T cells (Figure 2A). After these constructs were further packaged into lentiviral vectors, to compare the efficacy of safety switch side by side in both human pan-T and Jurkat cells, the lentiviral vectors expressing the engineered CAR constructs were transduced into both cell types at MOI=10. The expression of transduced CAR transgenes was confirmed by using flow cytometry via Protein L staining which binds to immunoglobulin kappa light chains. Lentiviral vector *hEF1α-CAR-tBID* had 13% transduction rate in human pan-T cells and 52% transduction rate in Jurkat cells (Supplementary Figure 4). Lentiviral vector *hEF1α-CAR-eGFP* had 17% transduction rate in human pan-T cells (Supplementary Figure 4). Lower transduction rate was observed in human pan-T cells for the same lentiviral vector.

**Figure 2.**
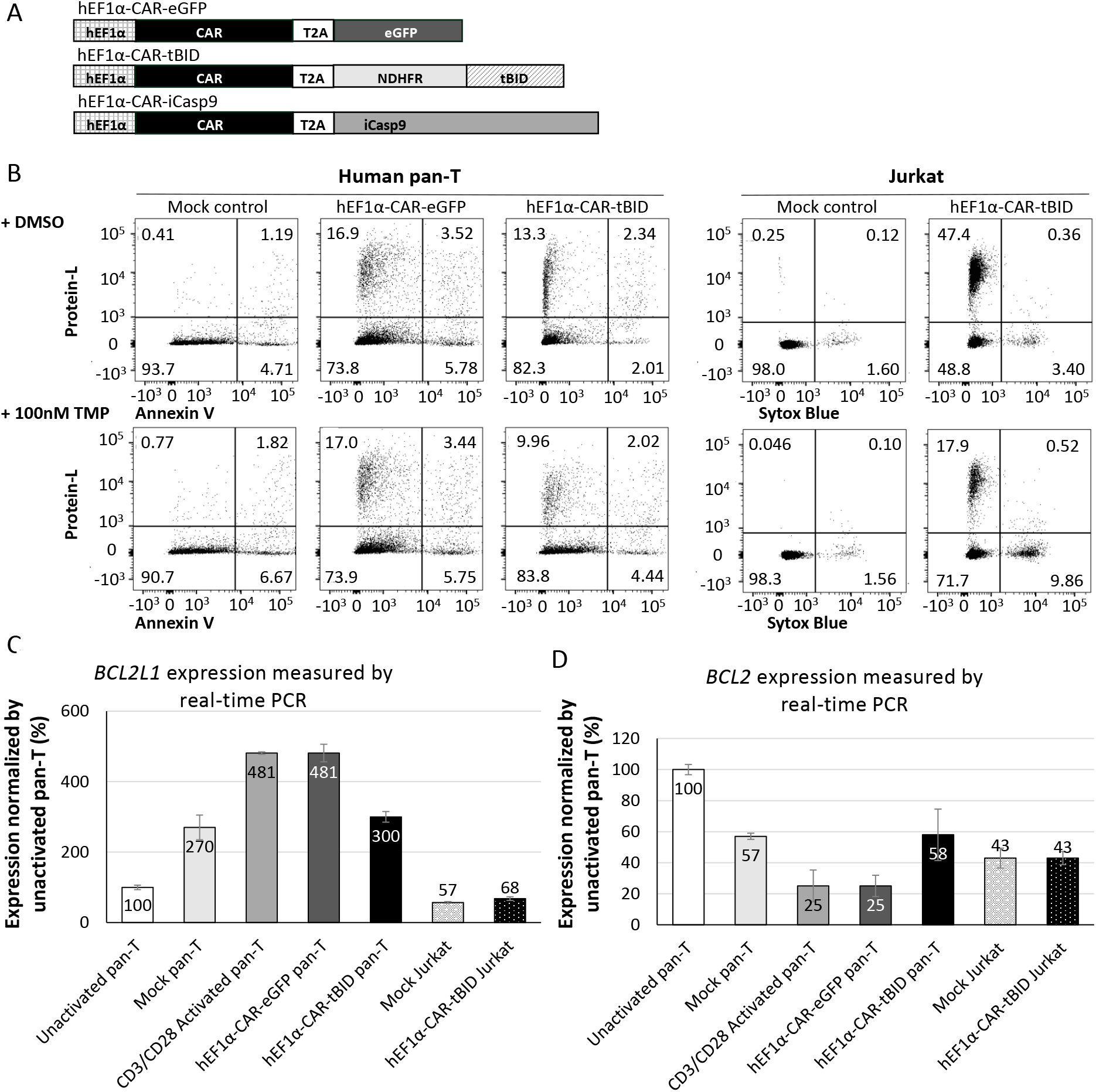
The killing efficacy of NDHFR-tBID safety switch in human pan-Tcells. A) Schematic diagrams of the CAR lentivirus vectors used to transduce human pan-T cells and Jurkat cells. B) Flow cytometry analysis of human pan-T cells and Jurkat cells transduced with CAR lentivirus vectors described in A at 72hr post DMSO or TMP treatment. All CAR lentivirus vectors described in A did not express any antibiotic selectable marker. Transduced human pan-T cells and Jurkat cells were treated with DMSO, or TMP on day 3 post lentivirus transduction. Cells were stained by Protein L and viability dye Annexin V or Sytox and analyzed by flow cytometry. C) Real-time PCR (TagMan) results for *BCL2L1* gene expression levels in human pan-T cells and Jurkat cells before and after activation or lentivirus transduction. D) Real-time PCR (TagMan) results for *BCL2* gene expression levels in human pan-T cells and Jurkat cells before and after activation or lentivirus transduction. Error bars represent standard deviation.

On day 7 post transduction, lentivirus transduced human pan-T cells, Jurkat cells, along with mock control cells (cells went through transduction procedure without adding lentivirus) for both cell types were treated with either DMSO or 100nM TMP to induce the apoptotic activity of NDHFR-tBID. The efficacy of eliminating CAR-positive cells was measured through Protein L staining and flow cytometry at 72hr post induction. As shown in Figure 2B, at 72hr after induction, the CAR-positive Jurkat cells that were transduced with *hEF1α-CAR-tBID* were reduced from 47.4% in DMSO treated group to 17.9% in TMP treated group, while no cytotoxicity was detected in the mock control group. This indicated strong killing efficacy of NDHFR-tBID in eliminating CAR positive Jurkat cells. However, the CAR-positive human pan-T cells transduced with *hEF1α-CAR-tBID* were only reduced from 13.3% in DMSO treated group to ~10% in TMP treated group during the same duration of treatment (Figure 2B). Therefore, NDHFR-tBID demonstrated much weaker killing potency in human pan-T cells than in Jurkat cells.

### BCL-XL and BCL-2 in human pan-T cells may inhibit the activity of NDHFR-tBID safety switch

As a death ligand, BID activates the mitochondrial membrane-bound receptor BAX through heterodimerizing the domain of BID with the BH1 domain of BAX (7). As a pro-apoptotic BCL-2 family member, BAX promotes cell death after being activated by BID through forming large oligomers that permeabilize the outer mitochondrial membrane, thereby committing cells to apoptosis. BCL-XL and BCL-2 are anti-apoptotic BCL-2 family members that prevent apoptosis induced by a variety of death stimuli (8,12,13). The BH1 domain of BCL-2 also interacts with the BH3 domain of BID (7), with heterodimerization between BID and BCL-2 inhibiting apoptosis (7). Previous studies have also shown that human T cells overexpressing BCL-2 were resistant to cell death (14). BAX also heterodimerizes with BCL-2 or BCL-XL(15). BCL-XL is structurally similar to BAX and competes with BAX for activation through interaction with membranes, BID, or BID activated BAX, thereby inhibiting BAX binding to membranes, oligomerization, and membrane permeabilization (16). It has been shown that anti-CD3 and anti-CD28 coactivation of T cells could strongly upregulate BCL-XL expression to promote survival of activated T cells (17,18). Since multiple lines of established studies have shown that BCL-XL and BCL-2 inhibits the apoptotic changes induced by tBID (8,13), we then hypothesize that BCL-2 and upregulated BCL-XL in activated human pan-T cells could contribute to the attenuation of NDHFR-tBID safety switch. To test this hypothesis, real-time PCR (TaqMan) was performed to validate the gene expression levels of *BCL2* (encodes BCL-2) and *BCL2L1* (encodes BCL-XL) in human pan-T cells and Jurkat cells pre- and post-transduction. As shown in Figure 2C, upon activation by anti-CD3/anti-CD28 Dynabeads or viral transduction, the *BCL2L1* expression was elevated by 3~5-fold in human pan-T cells. However, the *BCL2L1* expression was low and stable in Jurkat cells after transduction. After activation or transduction, human pan-T cells had 4~7-fold higher *BCL2L1* than Jurkat cells (Figure 2C). The *BCL2* expression was slightly higher (~1.4-fold) in human pan-T cells than in Jurkat cells after the transduction of *hEF1α-CAR-tBID* lentivirus (Figure 2D). High level of *BCL2L1* was detected when NDHFR-tBID activity was attenuated in activated human pan-T cells. Overall, our observation suggests that differential expression of pro- or anti-apoptotic pathway components between different cell types or preparations may affect the performance of our safety switch in the tested cells.

### Improving the potency of tBID safety switch in human pan-T cells through mutating in BH3 domain and fusing with CDHFR

Because elevated levels of BCL-XL and BCL-2 expression in human pan-T might result in the low efficacy of NDHFR-tBID safety switch, to address these issues, we inspected and fine-tuned the killing switch system by increasing tBID expression and disrupting the inhibitory influence of BCL-XL and BCL-2 in human pan-T cells.

The reported leakage effect of C-terminal ecDHFR destabilizing domain (CDHFR) mutant (N18T/A19V) (9) (Supplementary Figure 1) has hampered the initial attempt to fuse it with tBID. However the CDHFR mutant offered higher levels of fusion protein expression upon TMP induction when compared with NDHFR fusion protein (9) (Supplementary Figure 1). We then fused CDHFR to the C-terminus of tBID (tBID-CDHFR) with the expectation of using stronger tBID expression to overcome the inhibitory effects of BCL-XL and BCL-2 (Figure 3A). The published studies in recombinant BID proteins had identified a mutant (M97A/D98A) that specifically disrupted the interaction between BID and BCL-2 (2,7). We therefore incorporated the M97A/D98A mutation to the NDHFR-tBID construct, *hEF1α-CAR-tBID* (*M*), aiming to attenuate the inhibitory influence of BCL-2 on tBID safety switch (Figure 3A). Lentiviral constructs carrying these modifications were packaged (Figure 3A) and transduced into human pan-T cells at MOI=10. Due to the low transduction efficiency observed in human pan-T cells, the positively transduced cells expressing the CAR transgene were enriched by Protein L beads prior to further treatment (Supplementary Figure 5). To validate the killing efficacy of the modified tBID safety switches, the transduced human pan-T cells after Protein L beads enrichment were treated with either DMSO or 100nM TMP. The efficacy of eliminating CAR-positive cells was measured by flow cytometry via Protein L staining at 72hr post compound treatment (Figure 3B). As demonstrated in Figure 3B, addition of the M97A/D98A mutation to tBID (*hEF1α-CAR-tBID* (*M*)) slightly enhanced the killing potency of NDHFR-tBID by reducing CAR positive population from 75.3% (DMSO-treated) to 59.7% (TMP-treated). On the other hand, the non-modified NDHFR-tBID (*hEF1α-CAR-tBID*) only reduced CAR positive population from 79.1% (DMSO treated) to 71.1% (TMP treated). Fusing CDHFR with tBID provided improved potency as TMP treated *hEF1α-CAR-tBID-CDHFR* transduced cells reduced CAR-positive population to 38.8% compared with 77.2% in DMSO treated group. While analyzing the flow cytometry data, we noticed that the total viable cell number was significantly reduced in *hEF1α-CAR-tBID-CDHFR* transduced cells after TMP treatment. We then combined cell counting beads along with flow cytometry analysis to quantify the killing efficacy of modified tBID safety switches more precisely. As shown in Figure 3C, DMSO- or 100nM TMP-treated transduced cells were stained with Protein L and viability dye at various time points after treatment, then counted with cell counting beads through flow cytometry for the CAR positive live cell number. While the control vector *hEF1α-CAR-eGFP* transduced human pan-T cells had the same viable CAR positive cells as mock control, the non-modified NDHFR-tBID construct, *hEF1α-CAR-tBID*, transduced viable CAR positive cells were gradually removed with 97% viable CAR-positive cells on day 1, 60% viable CAR-positive cells on day 2, and only 43% viable CAR-positive cells detected on day 3 post TMP treatment. Incorporating M97A/D98A to tBID (*hEF1α-CAR-tBID* (*M*)) slightly improved killing efficacy with detection of 68% viable CAR-positive cells on day-1, 58% viable CAR-positive cells on day-2, and 38% viable CAR-positive cells on day-3 post TMP treatment. The most robust killing potency was observed in *hEF1α-CAR-tBID-CDHFR* transduced cells that 59% viable CAR positive cells on day-1, 19% viable CAR positive cells on day-2, and only 7% viable CAR positive cells on day-3 could be detected post TMP treatment. These data indicated that fusing CDHFR with tBID significantly enhanced the killing efficacy while adding M97A/D98A mutations to tBID only slightly improved the killing efficacy.

**Figure 3.**
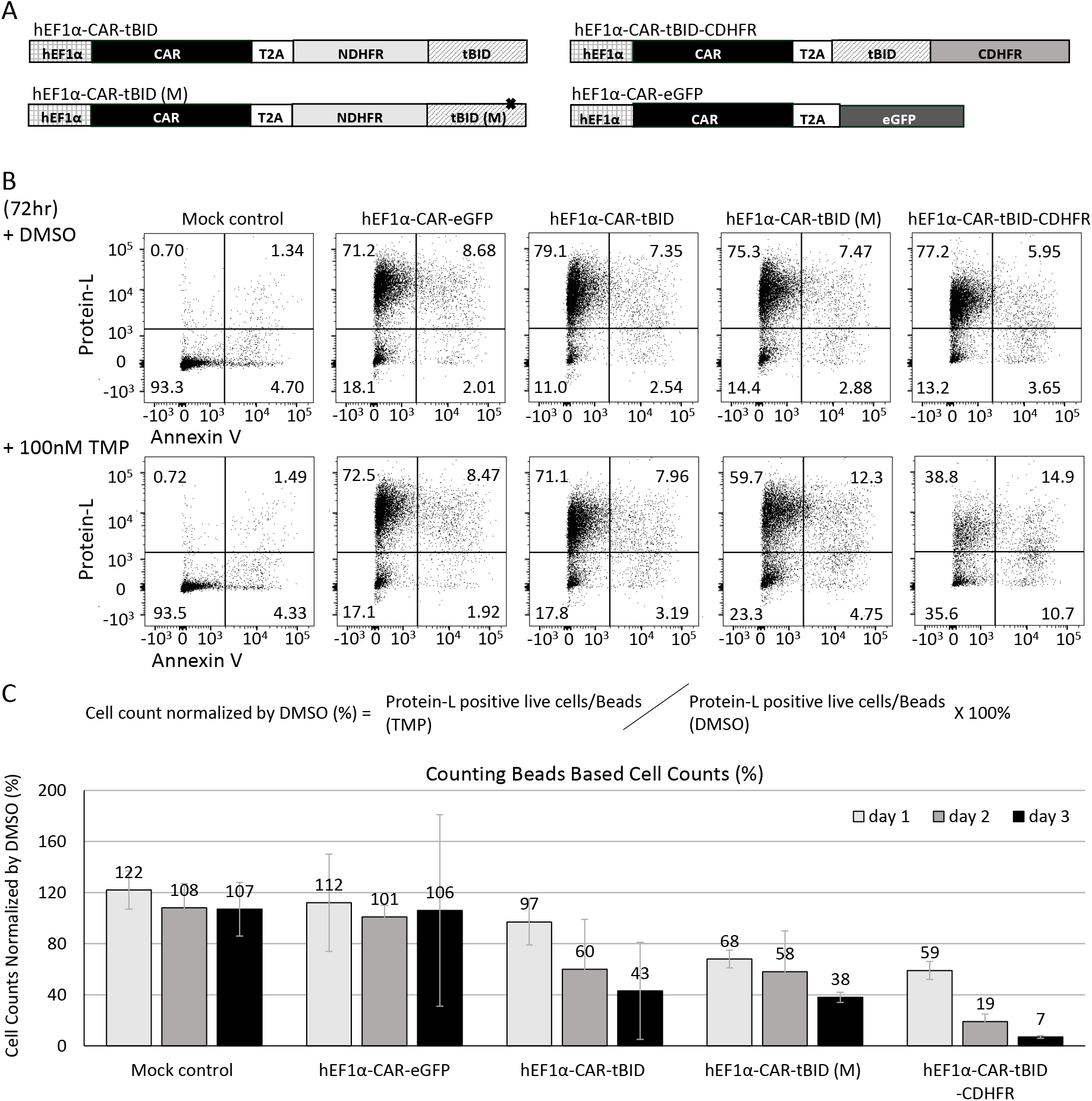
The killing efficacy of modified tBID safety switches in human pan-T cells. A) Schematic diagrams of the CAR lentivirus constructs with modified tBID safety switches used to transduce human pan-T cells. B) Flow cytometry analysis of human pan-T cells transduced with CAR lentivirus vectors described in A at 72hr post DMSO or TMP treatment. All CAR lentivirus vectors described in A did not express any antibiotic selectable marker. Transduced human pan-T cells were treated with DMSO or TMP on day 3 post lentivirus transduction. Cells were stained by Protein L and viability dye Annexin V and analyzed by flow cytometry. C) Viable cell count quantified by cell counting beads and flow cytometry analysis. Human pan-T cells transduced with lentivirus described in A were treated with DMSO or TMP on day 3 post transduction. The cells were stained with Protein L and Annexin V at various time points post treatment and quantified by cell counting beads during flow cytometry analysis. The cell counts in TMP treated groups were normalized by DMSO treat group and converted into percentage.

## DISCUSSION and CONCLUSION

In this study, we developed a safety switch based on the activity of tBID. In order to conveniently regulate the apoptotic activity of tBID, we took advantage of the cellular degradation machinery by fusing tBID to an engineered ecDHFR-derived destabilizing domain(9). The fusion of ecDHFR destabilizing domain to tBID confers the instability to entire fusion protein and causes rapid degradation. Application of ecDHFR specific inhibitor, TMP, stabilizes the destabilizing domain and the fusion tBID protein resulted in the activation of tBID apoptotic activity. The N-terminal destabilizing domain (mutant R12Y/Y100I) was first chosen to fuse with tBID in our study due to its high efficacy demonstrated by Mari Iwamoto et. al. (9). The high potency of this NDHFR-tBID safety switch was validated and proven in Jurkat cells. However, the NDHFR-tBID safety switch failed to induce cell death in human pan-T cells. BCL-XL and BCL-2 are BCL-2 family members that prevent apoptosis induced by a variety of death stimuli (8,12,13). In addition, the activation of T cells through anti-CD3 and anti-CD28 strongly upregulates BCL-XL expression to promote survival of activated T cells (17,18). In order to explore the reasons for NDHFR-tBID attenuation in human pan-T cells, we looked for potential inhibitors of tBID in human pan-T cells and potential methods to enhance the killing efficacy of tBID. The real-time PCR we performed demonstrated that human pan-T cells had 4~7-fold higher *BCL2L1* gene expression and slightly higher (1.4-fold) *BCL2* gene expression than Jurkat cells after activation or transduction. Therefore, both BCL-2 and BCL-XL could serve as inhibitors of tBID based safety switch in activated human pan-T cells. This limits the application of NDHFR-tBID in cells that have high levels of BCL-XL and BCL-2 expression. To overcome this limitation, we incorporated mutations (M97A/D98A) that have been demonstrated to disrupt the BCL-2 inhibition on tBID (2,7). The resulted modified NDHFR-tBID (M) safety switch did display stronger killing potency compared to nonmodified NDHFR-tBID (Figure 3B, 3C). However more striking improvement was achieved by switching the N-terminal destabilizing domain (mutant R12Y/Y100I) to the C-terminal destabilizing domain (mutant N18T/A19V) (9). C-terminal destabilizing domain fusion proteins have demonstrated higher expression levels upon TMP treatment than N-terminal destabilizing domain fusion proteins (Supplementary Figure 1) (9), however the leakage effects from C-terminal destabilizing domain hindered the initial attempt to fuse it with tBID in Jurkat cells (Supplementary Figure 1). The C-terminal destabilizing domain fused tBID safety switch tBID-CDHFR has demonstrated robust killing efficacy upon TMP treatment in human pan-T cells (Figure 3B, 3C), indicating stronger tBID expression could mitigate some inhibitory effects of BCL-XL and BCL-2. Considering the potential basal toxicity of the C-terminal destabilizing domain, we did not incorporate the M97A/D98A mutations to tBID-CDHFR safety switch. Although the potency of tBID based safety switch was improved after fusing tBID with CDHFR or incorporating M97A/D98A mutations under our *in vitro* condition, the inhibitory effects of BCL-XL and BCL-2 on tBID safety switch need to be considered when applying to other cell types.

The optimal safety switch will be fast-acting, with no or low immunogenicity, no or low basal toxicity and induced through a non-toxic small molecule. The tBID-CDHFR safety switch achieved a killing outcome between two clinically-validated safety switches, iCasp9 and HSV-TK, and enabled to kill ~90% transduced human pan-T cells in 3 days after 100nM TMP treatment in our *in vitro* assay (Figure 3C). The immunogenicity of ecDHFR-derived destabilizing domain will need to be determined through further *in vivo* studies and might potentially be mitigated through codon optimization.

Since we compared iCasp9 and tBID based safety switch in Jurkat cells, we planned to do the same side by side comparation of these safety switches in human pan-T cells as well. However, despite many attempts, we were not able to package *hEF1α-CAR-iCasp9* into enough lentiviral vector due to low viability of the HEK293 cells used to package this construct (Supplementary Table 1). There could be some element inhibiting lentiviral packaging in this vector which requires further modification or expression of *CAR-iCasp9* during lentiviral vector packaging was detrimental to HEK293 cell’s viability. We therefore removed this vector from testing list in human pan-T cells. No such toxicity was observed when packaging tBID constructs into lentiviral vector during this study. TMP as a very well-established antibiotic used to treat bacterial infections has been demonstrated to stabilize ecDHFR in a rapid, reversible and dose-dependent format. In summary we have developed a fast-acting tBID based safety switch with low basal toxicity that could be regulated through a commercially available, inexpensive, and blood-brain barrier permeable compound, TMP. Further investigation or characterization for applications of the tBID-based safety switch *in vivo* should be worthy to pursuit.

## ACKNOWLEDGEMENTS

The authors would like to thank Lili Yue and Qingwen Cheng for sharing experimental reagents.

## FOOTNOTE

### Consent for publication

All authors read and approved the final draft of the manuscript. This manuscript also has been reviewed and approved by Amgen Final Publication Review (FPR) process.

### Availability of data and materials

All data generated or analyzed during this study are included in this published article and its supplementary information files

### Author contributions

(I) Conception and design: J Lu, P Collins, S Wang; (II) Administrative support: CM Li, S Wang; (III) Provision of study materials or patients: J Lu, P Collins; (IV) Collection and assembly of data: J Lu; (V) Data analysis and interpretation: All authors; (VI) Manuscript writing: All authors; (VII) Final approval of manuscript: All authors

### Ethical Statement

Human T-cell samples were purchased anonymously from the vendor and considered exempt from IRB approval.

## ABBREVIATIONS & SYMBOLS

CRS: Cytokine Release Syndrome
tBID: truncated BID
HSV-TK: Herpes Simplex Virus Thymidine Kinase
iCasp9: Inducible Caspase 9
GCV: Ganciclovir
5-FC: 5-Fluorocytosine
CD: *E. coli*.-Derived Cytosine Deaminase
CID: Chemical Induction of Dimerization
CAR: Chimeric Antigen Receptor
ecDHFR: *E. coli* Dihydrofolate Reductase
TMP: Trimethoprim
BBB: Blood-Brain Barrier
BRB: Blood-Retinal Barrier
NDHFR: N-terminal ecDHFR Destabilizing Domain
CDHFR: C-terminal ecDHFR Destabilizing Domain
RLuc: Humanized Renilla Luciferase
7AAD: 7-Amino-Actinomycin D
MSCV: Murine Stem Cell Virus
EF1a: Elongation Factor 1 alpha
BID: BH3 Interacting-Domain Death Agonist
BCL-2: B-Cell Lymphoma 2
BCL-XL: B-Cell Lymphoma-Extra Large
BAX: Bcl-2-Associated X Protein
BH1: BCL-2 Homology domain 1
BH3: BCL-2 Homology domain 3
DMSO: Dimethyl Sulfoxide

## SUPPLEMENTARY APPENDIX

**Supplementary Figure 1.**
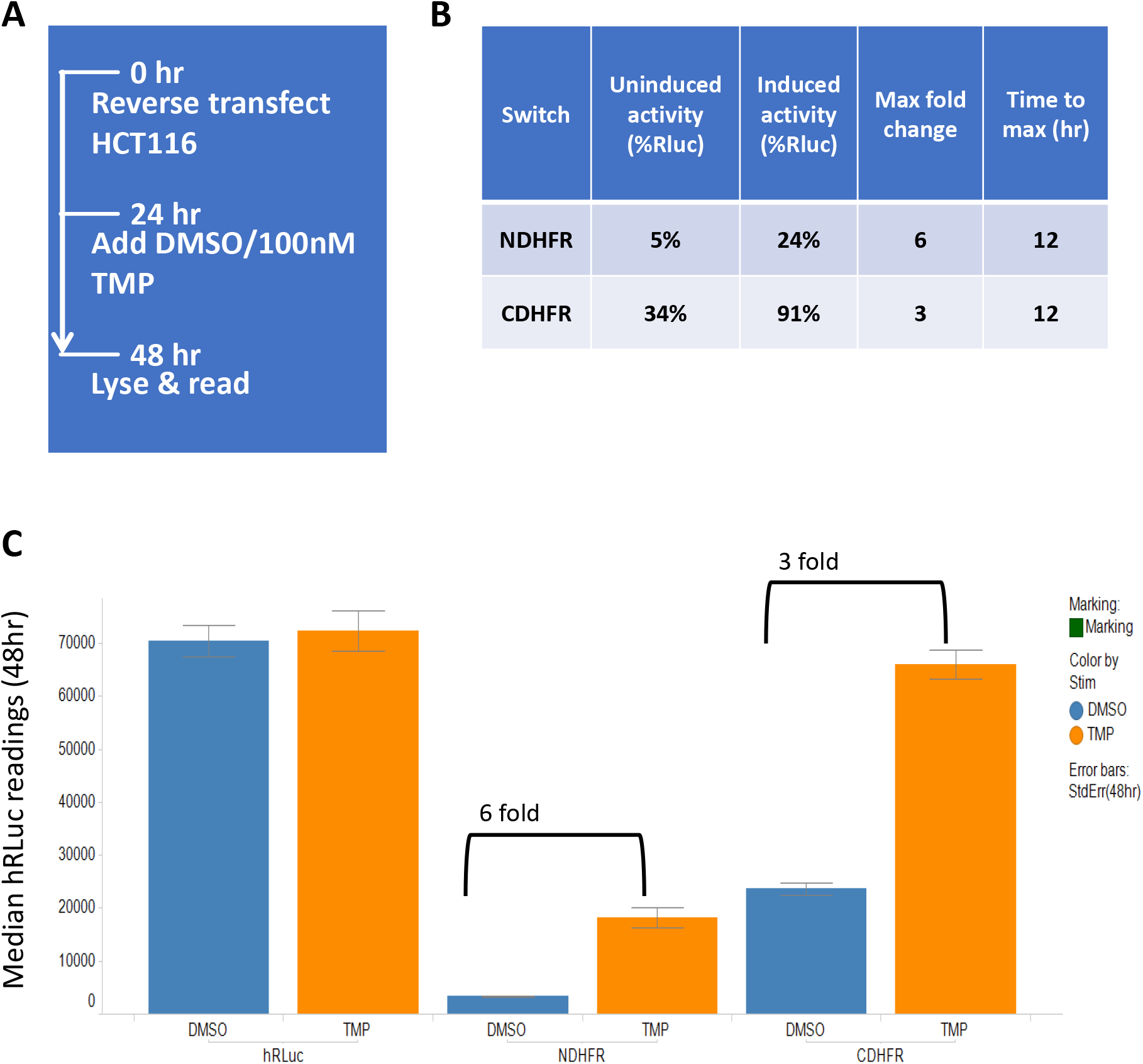
Compared the regulation efficacy and potency of NDHFR and CDHFR in HCT116 cells. A) Outline of the experiment procedure. B) Table to compare the efficacy and potency of NDHFR and CDHFR. C) Bar chart of hRLuc readings at 48hr post TMP or DMSO (vehicle) treatment.

**Supplementary Figure 2.**
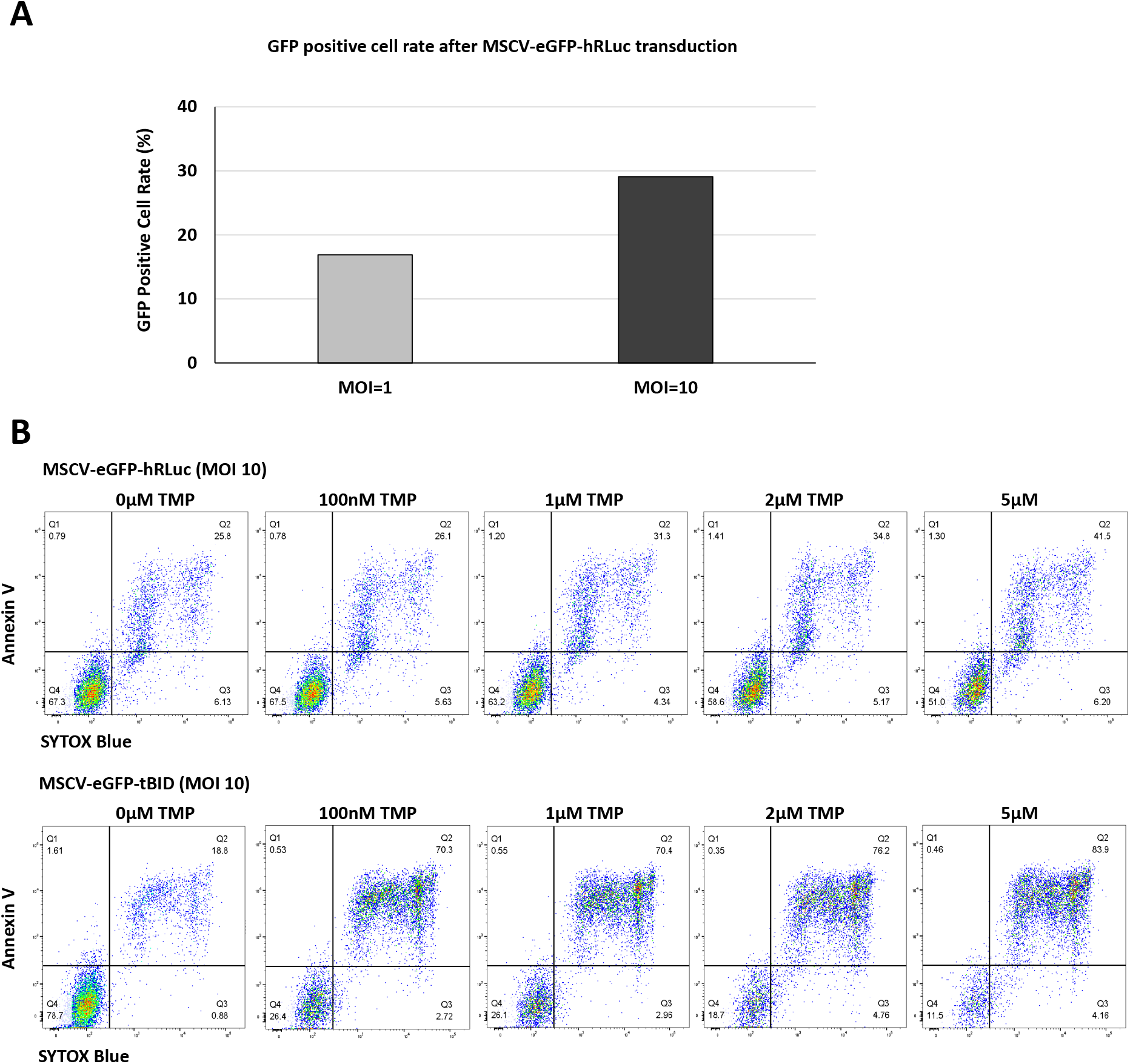
Optimization of lentivirus transduction MOI and TMP dosage. A) Comparison of lentivirus transduction efficacy in Jurkat cells at MOI 1 and MOI 10 by using *MSCV-eGFP-hRLuc*. The transduction efficacy was determined by the rate of GFP positive cells obtained from flow cytometry analysis. B) Dosage test of TMP for activating tBID safety switch in Jurkat cells. The Jurkat cells transduced by *MSCV-eGFP-hRLuc* lentivirus or *MSCV-eGFP-tBID* lentivirus were treated with TMP at numerous concentrations for two days. The viability of GFP positive transduced Jurkat cells were quantified through flow cytometry analysis after staining with Annexin V and SYTOX Blue.

**Supplementary Figure 3.**
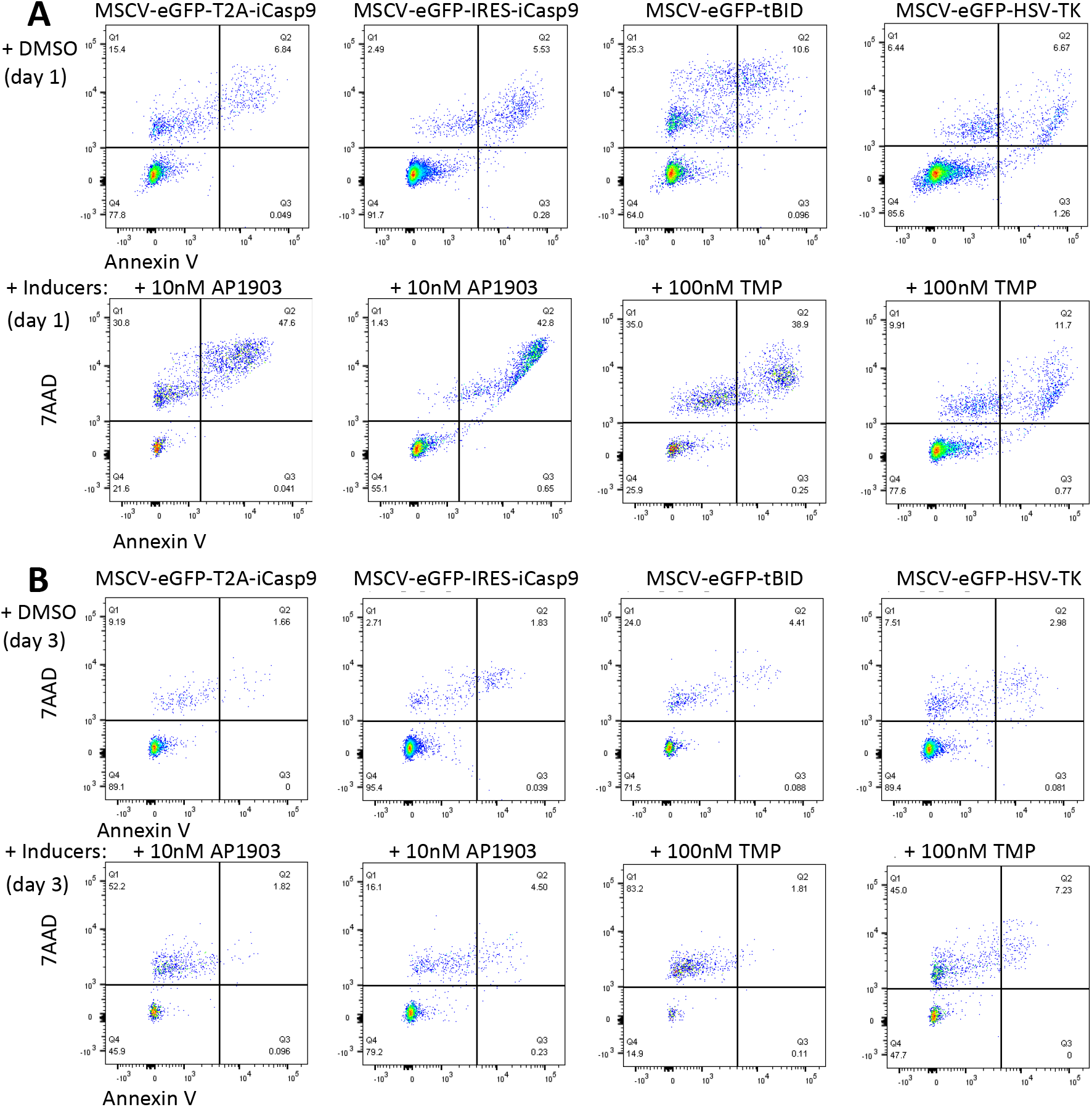
The killing efficacy of NDHFR-tBID safety switch in Jurkat cells. A) Flow cytometry analysis of Jurkat cells transduced with lentivirus vectors described in Figure 1A on day 1 post treatment. B) Flow cytometry analysis of Jurkat cells transduced with lentivirus vectors described in Figure 1A on day 3 post treatment.

**Supplementary Figure 4.**
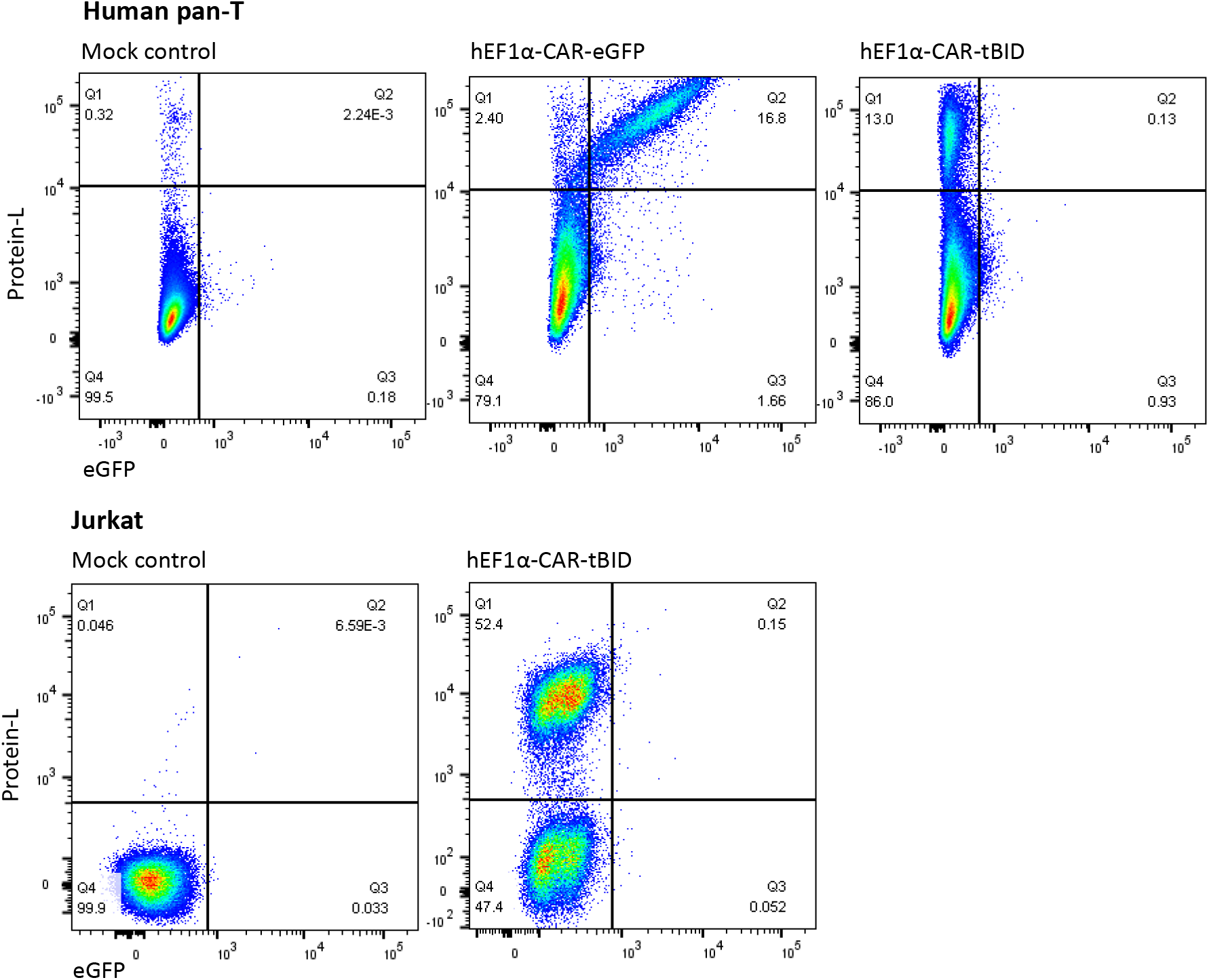
Transduction efficiency of lentiviral vectors in human pan-T cells and Jurkat cells. Lentiviral vectors *hEF1α-CAR-eGFP* and *hEF1α-CAR-tBID* were transduced into human pan-T cells at MOI=10. Lentiviral vector *hEF1α-CAR-tBID* was transduced into Jurkat cells at MOI=10. Both human pan-T cells and Jurkat cells were stained with Annexin V and Protein L followed by flow cytometry analysis on day 3 post transduction. The Annexin V negative single cells were further categorized for eGEP expression and positive Protain L staining.

**Supplementary Figure 5.**
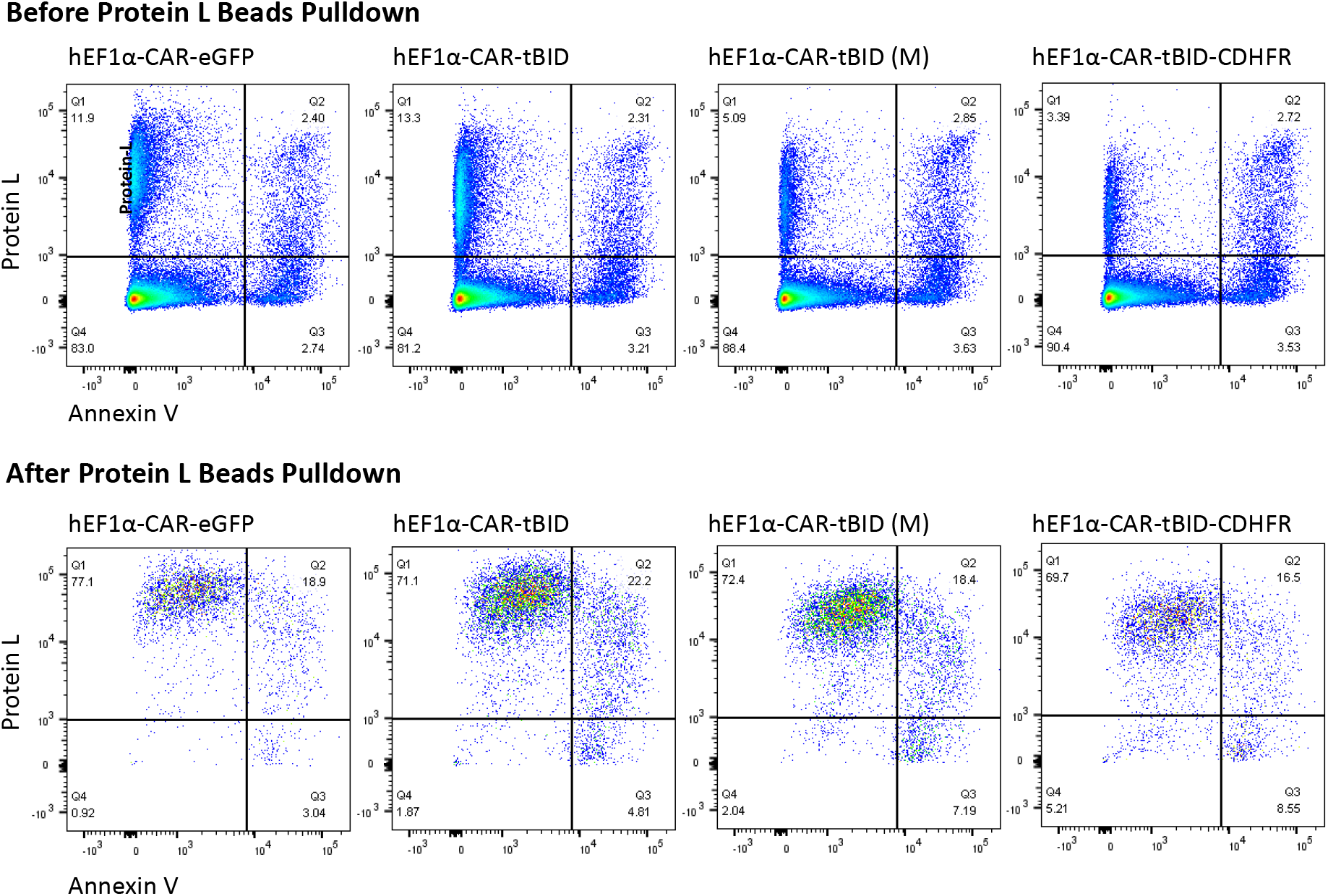
Protein L bead purification of lentiviral vector transduced human pan-T cells.

**Supplementary Table 1.** Comparison of HEK293 cell viability during lentivirus packaging

| Lentiviral Prep Name | In-HouseP24<br>(VP/mL) | In-House HEK293<br>Cell Viability (%)<br>Day of Harvest |
| --- | --- | --- |
| CAR-NDHFR-tBID | 4.63E+11 | 72.0 |
| CAR-iCasp9 | 8.80E+09 | 59.7 |
| CAR-eGFP | 5.43E+11 | 73.3 |

